# Global identification of mammalian host and nested gene pairs reveal tissue-specific transcriptional interplay

**DOI:** 10.1101/2023.05.02.539096

**Authors:** Bertille Montibus, James Cain, Rocio T Martinez-Nunez, Rebecca J. Oakey

## Abstract

Nucleotide sequences along a gene provide instructions to transcriptional and co-transcriptional machinery allowing genome expansion into the transcriptome. Interestingly, nucleotide sequence can often be shared between two genes and in some occurrences, a gene is located completely within a different gene, these are known as host/nested genes pairs. In these instances, if both genes are transcribed, overlap can result in a transcriptional crosstalk where genes regulate each other. Despite this, a comprehensive annotation of where such genes are located, and their expression patterns is lacking. To address this, we provide an up-to-date catalogue of host/nested gene pairs in mouse and human, showing that over a tenth of all genes contain a nested gene. We discovered that transcriptional co-occurrence is often tissue-specific. This co-expression was especially prevalent within the transcriptionally permissive tissue, testis. We used this developmental system and scRNA-seq analysis to demonstrate that co-expression of pairs can occur in single cells and transcription in the same place at the same time can enhance transcript diversity of the host gene. In agreement, host genes are more transcript diverse than the rest of the transcriptome and we propose that nested gene expression drives this observed diversity. Given that host/nested gene configurations were common in both human and mouse genomes, the interplay between pairs is therefore likely selected for, highlighting the relevance of transcriptional crosstalk between genes which share nucleic acid sequence. The results and analysis are available on an Rshiny application.

## INTRODUCTION

Classically genes are represented one after another along the chromosome’s genomic sequence separated by intergenic stretches of DNA which are devoid of genes, in a strand-specific manner. However, in practice, the genomic organisation is more complex, and genes can in reality occupy the same genomic space as each other and are considered as overlapping genes. The first overlapping pairs of genes were identified in the genomes of viruses (Weisbeek et al. 1977). Viruses are rapidly evolving organisms, with a small genome that must be packaged into a small capsid, and still maintain genetic novelty (Feiss et al. 1977; Wu et al. 2010). Thus, overlapping their genes allows them to deal with these constrains while enhancing their evolutionary stability because a mutation in an overlapping pair would affect more than one gene (Simon-Loriere et al. 2013). Higher order organisms, on the other hand, contain vast amount of intergenic space for genes to occupy, do not have the requirement of rapid evolution and yet, still contain genomic overlaps (Veeramachaneni et al. 2004). In eukaryotes, the first of these was identified in 1986 in Drosophila and mouse (Henikoff et al. 1986; Spencer et al. 1986; Williams and Fried 1986).

In 2019, a study in humans estimated that about 26% of the protein coding genes overlapped with at least one other protein coding gene (Chen et al. 2019). Most of the time, the overlap involved a non-coding sequence (5’UnTranslated Region or UTR, 3’UTR or intronic sequence) and sometimes only a few nucleotides were involved at the 3’ or 5’ end of the genes. However, larger genomic overlaps exist and can result in a gene being fully contained within a completely different gene. This configuration of genes contained one within the other is known as a host/nested gene pair. Genes organised this way are likely to cooperatively regulate one other because they occupy the same genomic space.

The steric hindrance observed when two RNA polymerases transcribe two different genes in the same genomic region (Billingsley et al. 2012) and transcriptional collision observed at convergent genes (Prescott and Proudfoot 2002) suggests that host and nested genes may prevent one another’s expression and may be mutually exclusively expressed. Furthermore, a repressive chromatin environment that prevents instances of intragenic spurious initiation of transcription is established in gene bodies by transcription itself (Latos et al. 2012; Neri et al. 2017). This is likely to repress nested gene expression when the host gene is expressed. On the other hand, nested gene expression has been shown to impact host gene pre-mRNA processing mechanisms, and, as a consequence, can generate host RNA isoforms with different stability and coding capacity (Licatalosi and Darnell 2010). This was shown for example at two independent imprinted loci, where the expression of a nested gene is associated with “premature” alternative polyadenylation of the host gene upstream of the nested gene whereas its silencing is associated with polyadenylation at the canonical 3’UTR (Cowley et al. 2012; Wood et al. 2008). This extends beyond the imprinted context and has been suggested to occur at other host genes harbouring nested *LINE1* retrotransposons, nested genes or putative intragenic promoters (Amante et al. 2020; Kaer et al. 2011). Growing evidence indicates that the short isoforms generated through “premature” polyadenylation are key for cell functions (Singh et al. 2018).

Thus, even though the role and the consequences of the host/nested gene organisation are starting to be uncovered, an updated genome-wide analysis is urgently needed. As such, characterisation and identification of all possible host/nested genes is crucial for the understanding of gene regulation both in terms of level of expression and isoform regulation. Previous compilations of host/nested genes are now outdated. Two previous analyses using NCBI Refseq and microarray data, at the time, identified 373 and 128 host/nested gene pairs in human (Assis et al. 2008; Yu et al. 2005). Not only were the number of pairs identified different between the two analyses, but conclusions regarding co-regulation of the pairs were different. Yu *et al,* demonstrated that most pairs were anti-correlated, suggesting mainly a mutually exclusive expression pattern whilst some showed a positive correlation and were expressed at the same time (Yu et al. 2005). Assis *et al* did not identify any significant correlation between the pairs, suggesting that host and nested genes are not influencing each other’s expression and that this organisation is “neutral” (Assis et al. 2008).

These are the most recent collations of host nested genes in human from 2005 and 2008 respectively (Assis et al. 2008; Yu et al. 2005). Genomic annotation is now much more comprehensive and advances in measurement technologies detect transcript isoforms in a more systematic way, as such, these previous lists underestimate the actual numbers (Frankish et al. 2019). Here, we identify and characterize the host/nested gene pairs in mouse and human using contemporary genomic annotations, RNA-sequencing and scRNA-seq data, and collate this analysis into an accessible shiny app for use by the scientific community (https://hngeneviewer.sites.er.kcl.ac.uk/hn_viewer/).

## RESULTS

### About a sixth of human (17%) and a tenth of mouse (12%) genes contain at least one nested gene

To obtain the most accurate and exhaustive list of host and nested genes in mouse and human we decided to use the GENCODE consortium genomic annotation. We used the “comprehensive” set which includes the highest number of transcripts associated with protein-coding genes, pseudogenes, long non-coding RNAs and small non-coding RNAs genes (Frankish et al. 2019). Transcripts with “To be Experimentally Confirmed” and “immunoglobulin (IG) variable chain or T-cell receptor (TR) genes” biotypes were removed. In addition, complex loci where alternative promoter or recombined segments are annotated with different gene names but cannot be considered as independent genes were also removed (Jia and Wu 2020; Jung et al. 2006; Strassburg et al. 2008) (see methods and Figure 1A) to ensure that a gene was defined as a validated transcriptional unit. Coordinate information for all the remaining genes were overlapped and the result filtered according to the extent of the overlap. Only pairs where all the transcripts of one gene were fully contained inside another transcript and where this involved transcripts from genes with different names were retained. When different host transcripts of the same genes were involved in a pair, the longest was selected. After filtering 7560 and 13088 host/nested genes pairs in mouse and human remained respectively (Supplemental_Table_S1 and Supplemental_Table_S2). This represented 4801 (∼12% of the genes) and 7661 (∼17% of the genes) genes with at least one nested gene in mouse and human respectively (Figure 1A, Supplemental_Table_S1 and Supplemental_Table_S2)

**Figure 1.**
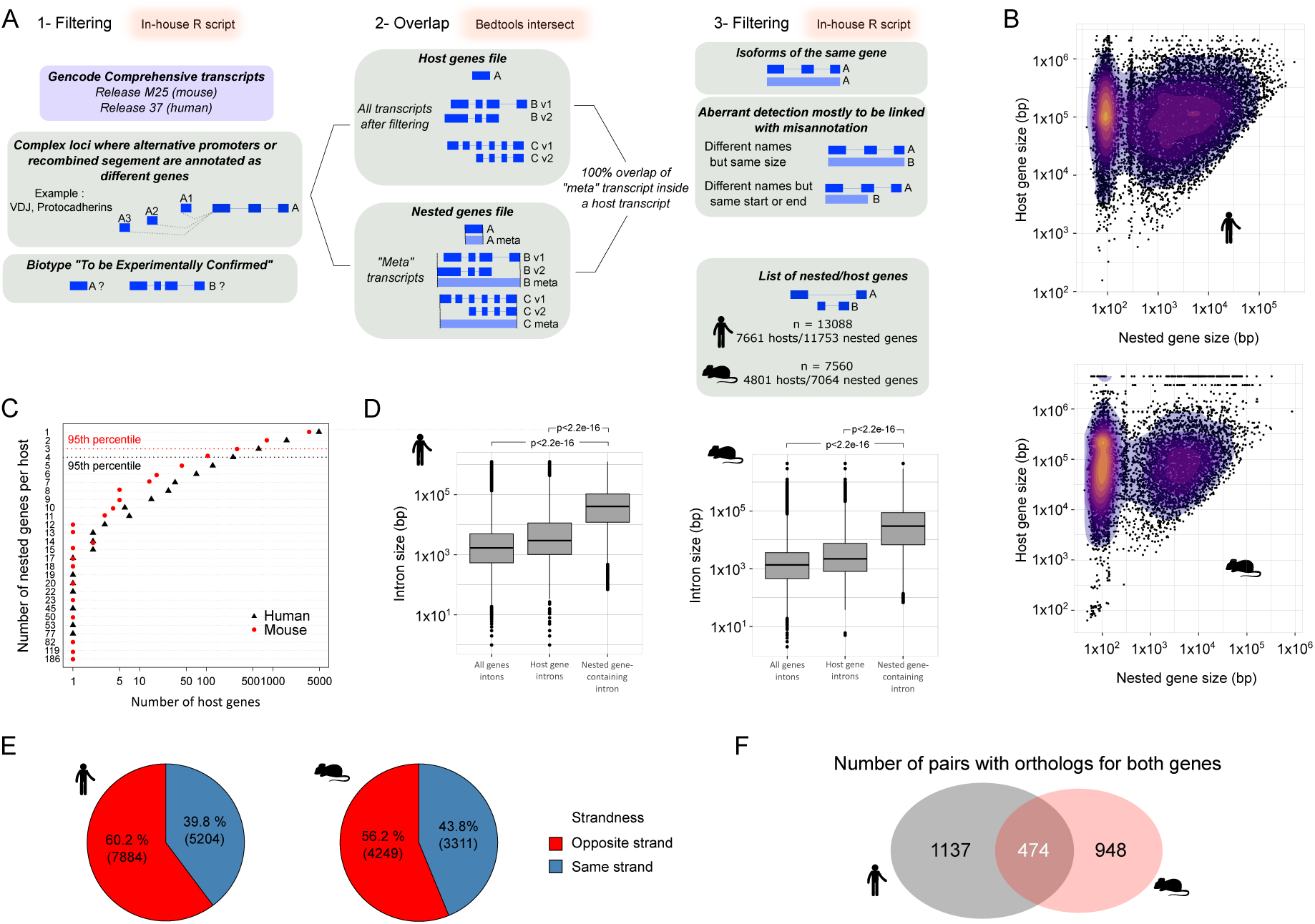
Host/nested genes pairs in mouse and human shows similar characteristics. (A) Schematic of the pipeline used to generate the list of host/nested genes pairs. (B) Scatter plot showing the correspondence between the sizes of host and nested genes. The colour gradient indicates the density. A base-10 log scale was used for the gene size. (C) Scatter plots showing the number of nested genes per host. The dotted line represents the 95th percentile of the distribution. (D) Distribution of the intron size for all introns in all the genes, all introns in host genes or the intron of the host containing a nested gene. A base-10 log scale was used for the intron size. The significance of the difference was tested using a Welch Two Sample t-test. The resulting p-value is mentioned on the graph. (E) Proportion of host/nested genes pairs with both genes in the same orientation or in the opposition direction. (F) Overlap and conservation between the pairs having an ortholog for both host and nested in the other species.

### Hosts harbour nested genes preferentially inside a long intron and show no strong orientation bias

To identify any preferences of host/nested gene structure and location, we performed a suite of analyses based on their basic characteristics. Unsurprisingly, host genes were longer than their nested gene partner and longer than the average size of the corresponding transcriptome (Supplemental_Fig_S1A). However, there was no correlation between the size of the host and their nested gene (Figure 1B). To minimize the impact to the transcriptome, we hypothesised that host genes would not contain a large number of nested genes and that they would preferentially locate within non-coding intronic host gene regions. Most of the host genes, indeed, harboured less than 4 nested genes (95th percentile, 3 in mouse and 4 in human) but in some extreme cases host genes could contain up to 77 nested genes in human and 186 in mouse (Figure 1C). In addition, ∼75% of the nested genes are fully contained inside one intron of their host gene and ∼20% of nested genes span both an intron and an exon (Supplemental_Fig_S1B). Surprisingly, the remaining overlap with exonic sequences and 3-5% of nested genes are completely embedded within exons (Supplemental_Fig_S1B). In this case however, nested genes predominantly localised to the last exon of host genes (Supplemental_Fig_S1D). Within introns, nested genes were often located in the first intron (Supplemental_Fig_S1C) and preferentially inside the largest host intron (Figure 1D), as observed before (Yu et al. 2005). Assessment of pair orientation did reveal a slight bias towards being in the opposite orientation between pairs, but the distribution between opposing and same orientation were broadly similar (Figure 1E). This pattern was similar for nested genes fully contained in introns and a stronger bias for opposite orientation was observed when nested genes spanned both exons and introns (Supplemental_Fig_S1B, Supplemental_Fig_S1C and Supplemental_Fig_S1D).

Given that host/nested gene pairs had a surprising coverage of over a tenth of genes across the genome, we asked if these pairs were conserved between mouse and human. Only pairs where both genes have an identified orthologue in the other species were considered. Using these filtered pairs, it was found that 29% of the human pairs and 33% of the mouse pairs were conserved in the other species (Figure 1F, Supplemental_Table_S3), as exemplified by *Mcph1*/*Angpt2* and *MCPH1*/*ANGPT2* (Supplemental_Fig_S1E).

### Host and Nested genes are enriched for different functional classes

To better characterise the genes involved in host/nested gene pairs, we took advantage of the broad functional classes of biotypes available in GENCODE (Frankish et al. 2019). By comparison to the distribution of biotypes among all the genes, we found that host genes were enriched for protein coding genes (p<0.001, hypergeometric test, Figure 2A). The nested genes were enriched for small RNAs in general whereas host genes were depleted for these transcripts in both species (p<0.001, hypergeometric test, Figure 2A). When looking at the number of instances for each biotype, we observed that most of the host genes, but also a high proportion of the nested genes were protein coding genes. As expected from the enrichments, we also detected a high number of non-coding RNA in the nested genes list (lncRNAs, antisense RNAs and miRNAs) (Supplemental_Fig_S2A). Finally, when considering the correspondence of biotypes between the host and nested genes per pair, no trend was observed for host and nested genes biotype association, other than the expected higher proportion of pairs of genes being organised in the opposite direction when one of the partners was annotated as an antisense transcript (Figure 2C).

**Figure 2.**
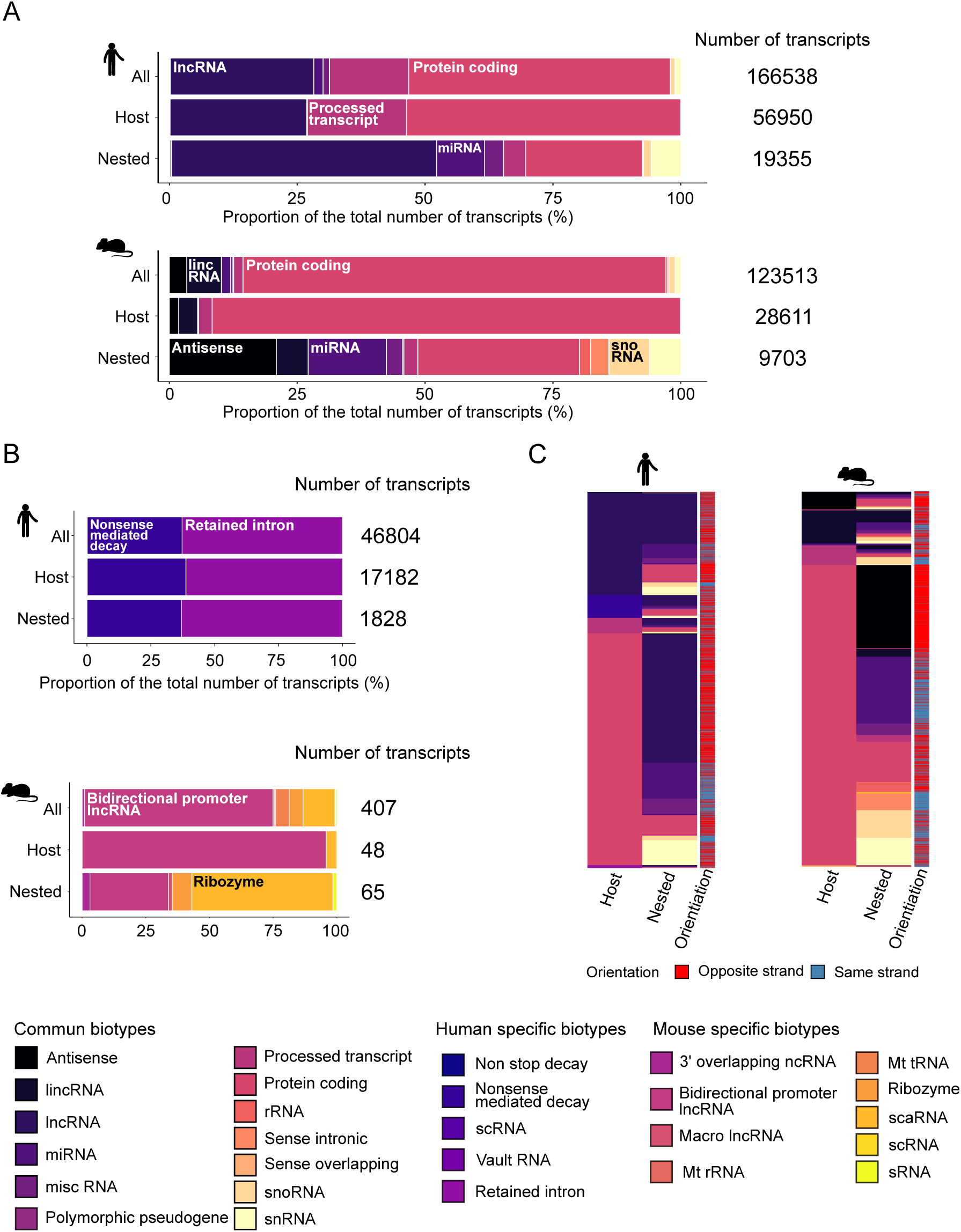
Host and nested gene biotypes. (A) Bar chart representing the proportions of biotype which are common across both species. Proportions are displayed for all genes, host genes and nested genes in human and mouse genomes. (B) Bar chart representing proportions of biotypes unique to human and mouse genome across all genes, host genes and nested genes. (C) Heatmap showing the correspondence between host and nested gene biotypes per pair and their orientation to each other.

### Host or Nested gene expression is not linked to tissue-specificity

Host and nested genes were enriched for distinct functional classes and so we sought to annotate their biological function via gene ontology and expression analysis. Ontology analysis shows that host genes are enriched for terms associated with neural differentiation and synapse development (Supplemental_Fig_S2B) and nested genes for terms associated with negative regulation of translation and gene silencing as expected given that a large proportion of nested genes were small RNAs (Supplemental_Fig_S2C).

The gene ontology analysis of host genes suggested that they could be enriched for genes important for tissue specification (Supplemental_Fig_S2B). Therefore, we decided to investigate whether the host genes were expressed in a tissue-specific way using bulk tissue ENCODE RNA-sequencing data (Davis et al. 2018) (Supplemental_Table_S5, Supplemental_Table_S5). As a first approach, we calculated the standard deviation of host and nested gene expression. It showed that host genes had a greater variance in expression level (Supplemental_Fig_S3A), suggesting that these genes are expressed in a limited set of tissues. As the standard deviation is influenced by the level of expression, we investigated the global level of expression across each tissue for host and nested genes in comparison to all genes. Higher expression level of the host genes and lower expression of the nested genes were observed for protein coding genes and some other biotypes (Figure 3A). The higher expression of host protein-coding genes was detected in all tissues tested, whereas nested protein-coding genes had lower expression levels (Supplemental_Fig_S3B and Supplemental_Fig_S3C). This suggests that the differences between standard deviations could be due to differences in basal expression levels. To explore this, we computed the τ (Tau) index which is a tissue-specificity metric (Yanai et al. 2005). The value of τ varies between 0 for housekeeping genes (or broadly expressed genes) and 1 for tissue-specific genes. In neither mouse nor human, was there a major difference in the distribution of the value of τ indicating that host and nested genes are not enriched for tissue-specific genes when their expression is considered separately (Figure 3B).

**Figure 3.**
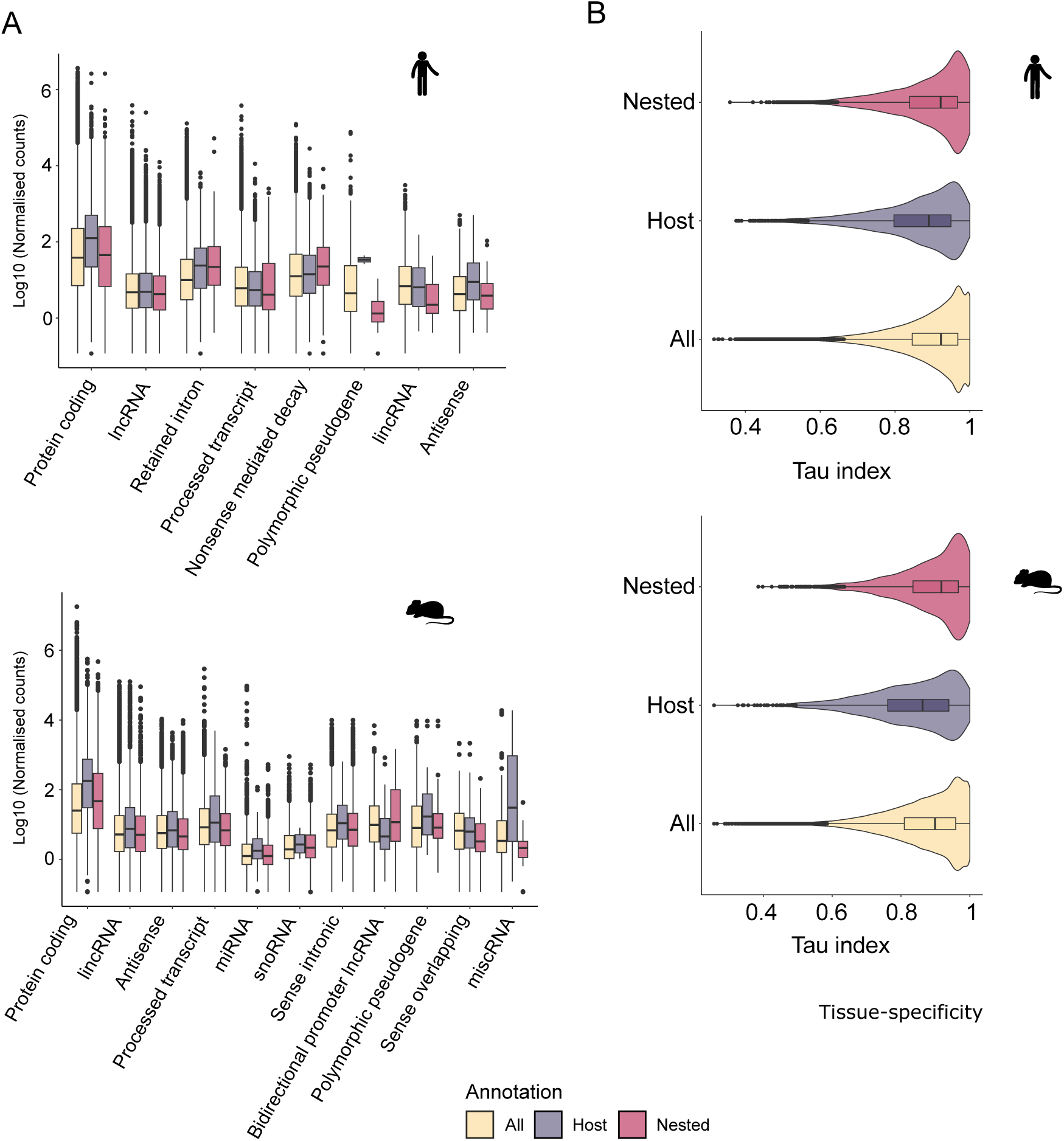
Host and nested gene expression profiles are not associated with tissue-specific expression. (A) Distribution of the expression of all, host, and nested genes for the biotypes associated with the three categories in human and mouse transcriptomes (B) Distribution of Tau (τ) index calculated using the normalised expression level detected in RNA-sequencing datasets. The higher the value of Tau the more the gene exhibit tissue specific expression.

### Co-expression of host and nested gene is linked to tissue-specificity

As intragenic elements have been previously associated with tissue-specificity (Amante et al. 2020), we hypothesised that co-expression could influence tissue-specific profiles and tested this by focusing on the correlation between host and nested genes expression. This also tested the model whereby there is a steric hindrance between two RNA polymerases transcribing in the same genomic region (Billingsley et al. 2012) and host gene expression represses nested gene expression (Neri et al. 2017). In this case, host and nested genes should be expressed in a mutually exclusive way. A Spearman’s rank correlation coefficient (ρ) was calculated for each pair between the expression profile of the host and the nested gene across tissues (as exemplified for the conserved pair *Mcph1*/*Angpt2* in Figure 4A). The distribution of ρ showed a peak between 0 and 0.5, indicating that, for most pairs, there is no positive or negative relationship between the expression level of the host and the nested genes in both species. This distribution was independent of host/nested gene pair orientation (Figure 4B, Supplemental_Fig_S4A).

**Figure 4.**
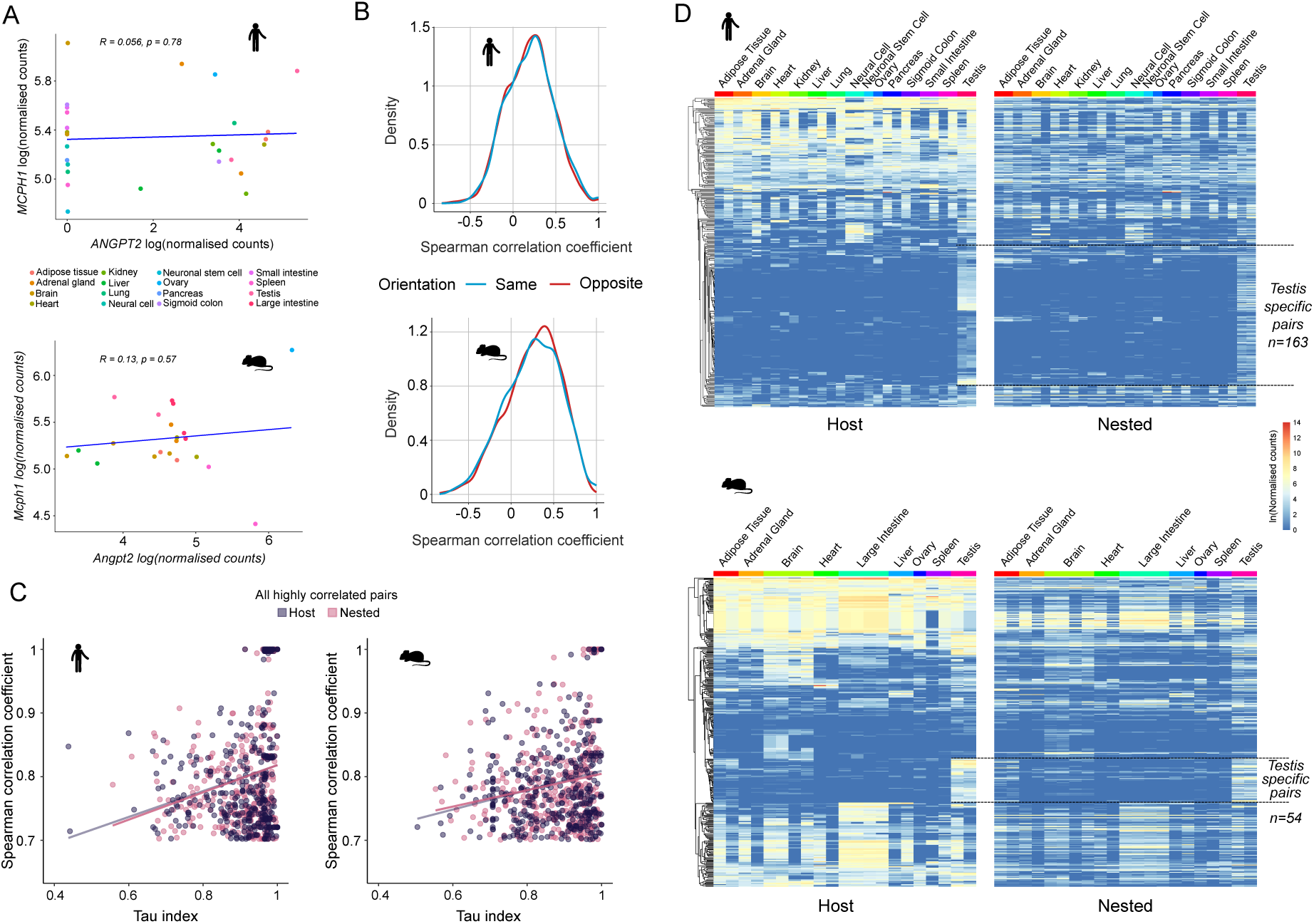
Tissue specific co-expression of host/nested gene pairs. (A) Examples of correlation plots used to calculate the Spearman’s rank correlation coefficient for the conserved pair *Mcph1*/*Angpt2*. (B) Distribution of the Spearman’s rank correlation coefficient for all host/nested genes pairs depending on their orientation to each other. (C) Correlation between the Spearman’s rank correlation coefficient value and the τ index for all the pairs exhibiting a Spearman’s rank correlation coefficient higher than 0.7. (D) Heatmap showing the normalised expression level detected in RNA-sequencing datasets for host and nested genes pairs with a Spearman’s rank correlation coefficient higher than 0.7. Host genes were clustered using hierarchical clustering according to their expression profile and their nested counterpart were maintained in the same order. The dotted lines are flanking the groups of pairs specifically co-expressed in testis.

In addition, anti-correlated expression was rare (ρ = <-0.7), whilst highly correlated pairs were more common (ρ = > 0.7) (Figure 4B). To determine whether this correlation was linked to the host and nested genes expression profile, the relationship between the correlation and the tissue-specificity was evaluated. The higher the correlation, the more the host and nested genes were expressed in a tissue specific manner as determined by the τ (Figure 4C, Supplemental_Fig_S4B). Examination of the expression profile of these highly correlated paris across tissues reveals a large group of pairs with co-expression specific to the testis (163 pairs, Figure 4D). This was in line with previous report of the testis showing both widespread gene activity and high transcriptome diversity (Soumillon et al. 2013). In addition vast changes in gene expression occur, in the testis during the process of spermatogenesis (Jan et al. 2017). Thus, the testis is both an excellent developmental and mechanistic model tissue to begin to understand the functional role of host and nested co-expression.

### Single-cell RNA sequencing reveals patterns of expression of host/nested genes pairs

Because, testis is a complex tissue, consisting of both somatic and germline cells at different stages of differentiation, bulk RNA-sequencing does not offer the necessary resolution to determine whether pairs of genes are expressed in the same cells or in different cells or are mutually exclusive. To assess this, a human testis single cell RNA-sequencing (scRNA-seq) dataset was leveraged (Di Persio et al. 2021). Expression of 64 out of 163 testis-specific pairs was detected in this dataset, which we attributed to the limited depth of single cell sequencing approaches. A Spearman’s rank correlation analysis was performed between host and nested gene expression per cell on the subset of testis-specific pairs identified by the ENCODE analysis (Figure 4D). Most of the pairs demonstrated a low absolute value of the Spearman’s rank correlation coefficient (Supplemental_Fig_S5A). However, as Spearman’s rank correlation is skewed by the low and zero counts of scRNA-seq data, it was not suitable to assess co-expression, nor, mutually exclusive expression of genes (Li and Li 2021; Pollen et al. 2014; Sanchez-Taltavull et al. 2020). Indeed, host/nested gene pairs demonstrate different profiles of co-expression despite similar Spearman’s rank correlation coefficients. (*AL109954.1/CST8*, R=0.29 and *RNASE11/AL163195.2*, R=0.25, Supplemental_Fig_S5B). To overcome this, we classified the single cells according to the host and nested gene expression based on a quarter of the maximum expression value for each gene (Supplemental_Fig_S5C). Expression above the threshold was indicative of robust detection and allows cell classification. Because we wanted to investigate the relationship between host and nested genes, we applied a filter of a minimum of 1% of the cells of the dataset had to have robust expression for both the host and the nested gene, selecting 34 pairs. Finally, expression above the threshold for both the host and the nested gene allows identification of cells which co-express the pair. Expression above one threshold but below the other indicates expression of only one member of the pair whereas expression below both thresholds classified the cells as non-expressing the pair (Supplemental_Fig_S5C). This generated an accurate representation of mutual exclusive expression and co-expression, e.g., for *LINC02253*/*AC020704.1* (0.06% of cells where both genes are expressed but 2.13% and 4.32% of cells with exclusive expression of the host and the nested gene, respectively) and *CASC16*/*AC026462.3* (16.64% of cells where both genes are expressed) respectively (Figure 5A).

**Figure 5.**
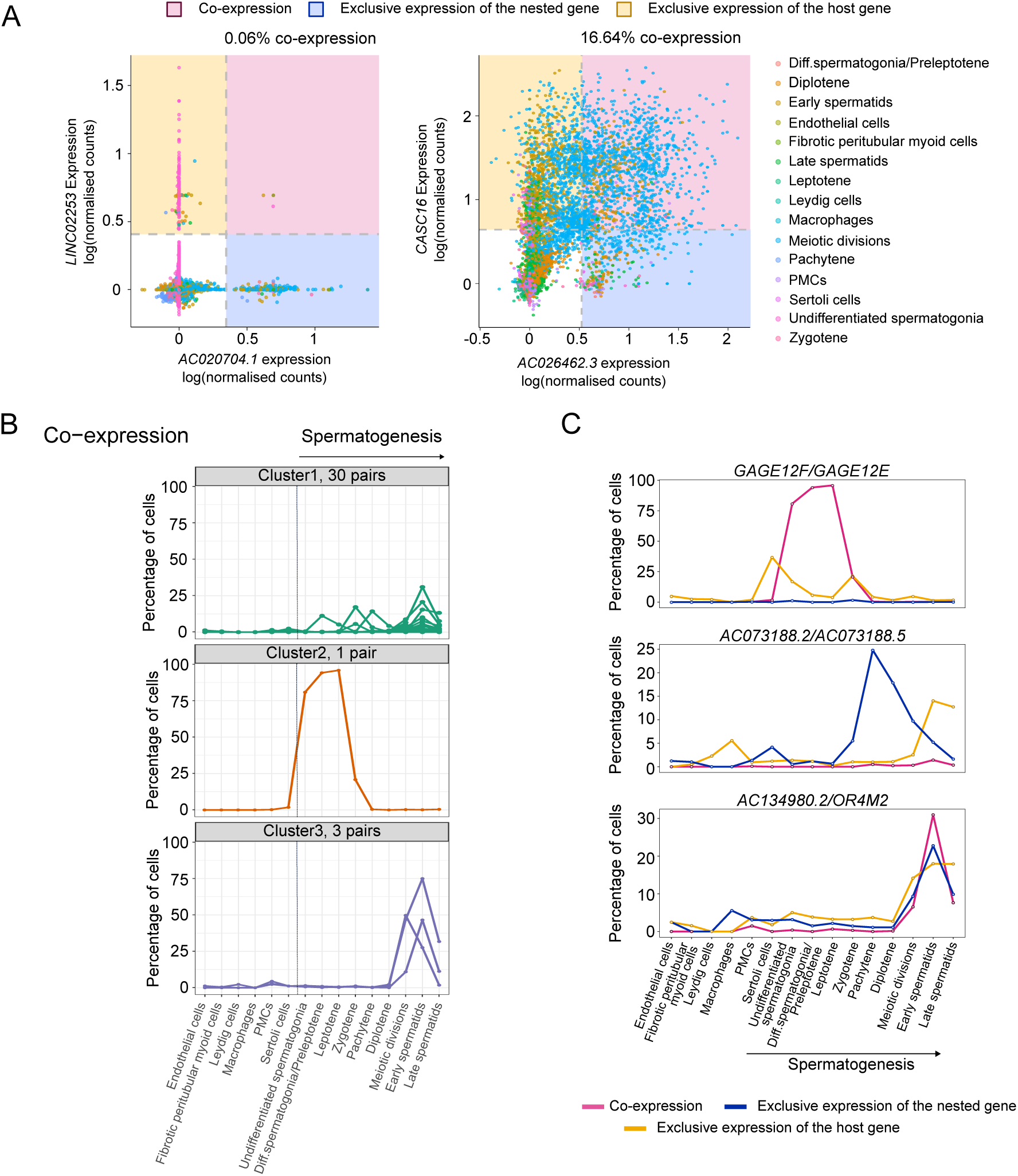
Host and nested genes with high co-expression in testis can exhibit dynamic co-expression patterns during spermatogenesis at the single cell level. (A) Scatter plots showing host and nested genes normalised expression level in testis single cells for two examples pairs: *LINC02253*/*AC020704.1* and *CASC16*/*AC026462.3*. (B) Proportion of cells which co-express host and nested genes pairs across spermatogenesis. Three distinct co-expression profiles were determined by K-means clustering. (C) Scatter plot showing the profiles of expression determined as in Supplemental_Fig_5C for three different pairs of host and nested genes. *CAGE12F/CAGE12E* is an example of pair where host and nested genes are mainly co-expressed, *AC073188.2/AC073188.5* an example of pair with mutually exclusive expression and *AC1349980.2/OR4M2* a pair with a complex expression profile in testis.

### Transcriptional interplay between host and nested genes during spermatogenesis

Co-expression or exclusive expression was limited to some of the cell types, as such we hypothesised that they are tightly regulated during development. For example, *AL163195.3*/*AL163195.2* was co-expressed mainly during the late stages of spermatogenesis (meiotic division, spermatids) and expression of *AL163195.3* alone was also observed at earlier stages (spermatogonia, leptotene, zygotene, pachytene) and in some of the somatic cells (Supplemental_Fig_S5D). Globally, the interplay between host and nested genes expression was defined for each pair based on the percentage of cells defined as co-expressing or mutually expressing in every cell type (Supplemental_Fig_S5E). We performed K-mean clustering (Figure 5B and Supplemental_Fig_S6A) and identified distinct profiles of co-expression or exclusive expression, indicating that host/nested gene expression and interplay is regulated according to cell types and during the process of spermatogenesis. However, a profile with a simple relationship (*i.e.* mostly co-expressed or host expressed in some cell types and nested gene in others) was rare. For example, *GAGE12E*/*GAGE12F* were mainly co-expressed and *AC073188.2/AC073188.5* exhibited mutually exclusive expression (Figure 5C). Most of the pairs showed a complex profile sometimes with co-expression and sometimes exclusive expression, as exemplified by *AC134980.2/OR4M2 and IGSF11/IGSF11-AS1* (Figure 5C and Supplemental_Fig_S6B). For some pairs presenting this more complex expression profile, we observed that while the host gene was broadly expressed, the nested gene expression was limited to testis (Figure 6D). Interestingly, *IGSF11* is a known regulator of meiosis during spermatogenesis (Chen et al. 2021) suggesting that the co-expression of host and nested genes and the changes in isoforms expression could be key for its function in testis.

**Figure 6.**
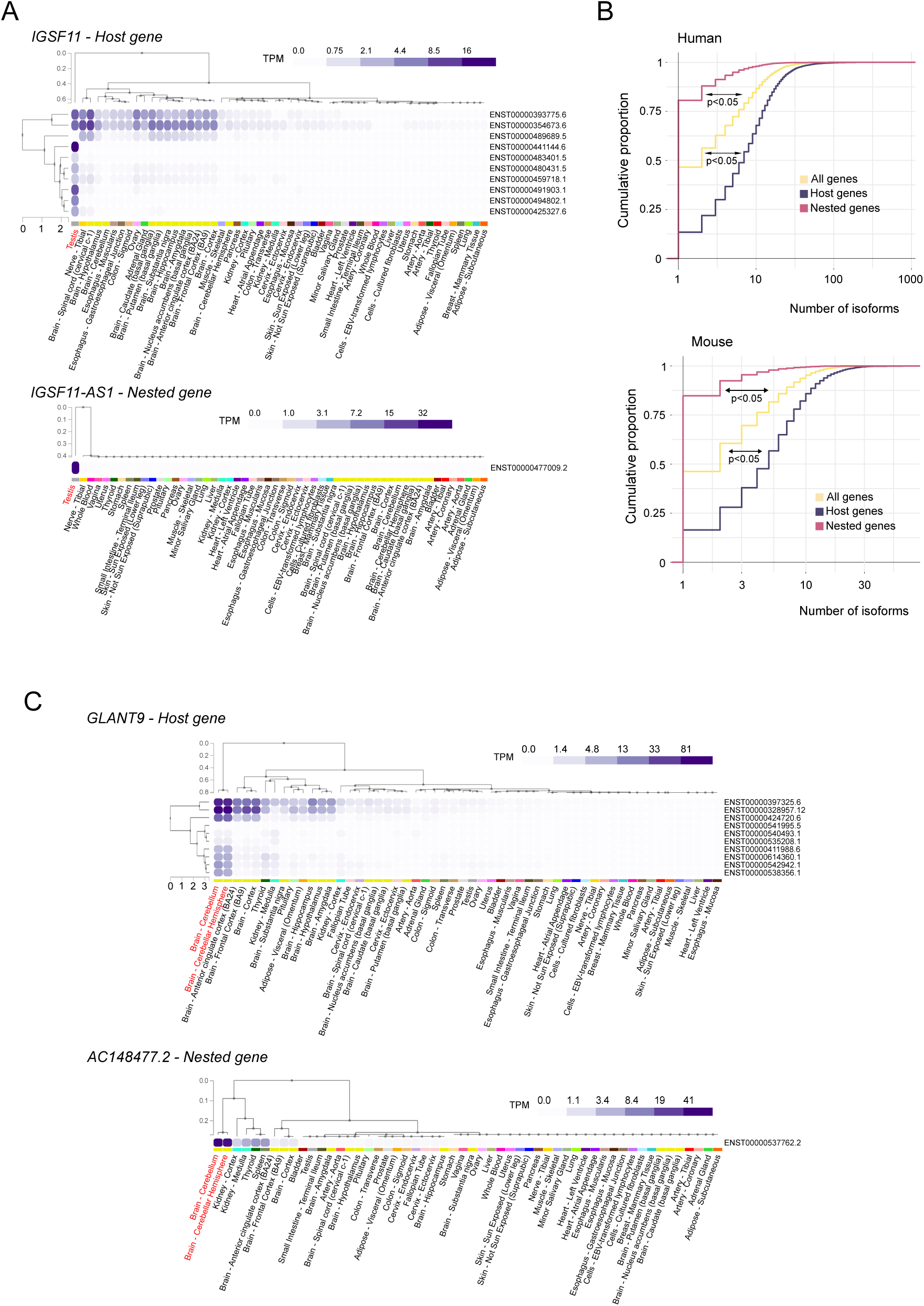
Host and nested genes co-expression is correlated with regulation of isoform diversity. (A) Profile of expression of the different isoforms (rows) of the *IGSF11/IGSF11-AS1* pair in different tissues (columns) from the GTEx portal data on isoforms expression (https://gtexportal.org/home/). Isoform and tissue were ordered by hierarchal clustering using the Euclidean distance and average linkages. Tissues where co-expression is happening are in red. (B) Cumulative proportion of the number of isoforms for host genes, nested genes and all genes. The significance of the difference was tested using a DTS statistic test to compare the empirical cumulative distribution functions (ECDF) (Dowd 2020). The resulting p-values are mentioned on the graph. (C) Profile of expression of the different isoforms (rows) of the *GLANT9/AC148477.2* pair in different tissues (columns) from the GTEx portal data on isoforms expression (https://gtexportal.org/home/). Isoform and tissue were ordered by hierarchal clustering using the Euclidean distance and average linkages. Tissues where co-expression is happening are in red.

### Host genes show a greater number of isoforms and co-expression correlates with changes in transcripts diversity

Using the GTEx portal data on isoform expression (Lonsdale et al. 2013), we observed that the nested gene expression restricted to testis was associated for this pair with a greater diversity of host isoforms expressed in testis (Figure 4D and Supplemental_Fig_S4B). The correlation between host specific isoforms and expression of the nested gene in testis was also observed for pairs which were not detected in the single cell RNA-seq dataset such as *MGAM/OR9A4* and *HMX1/AC116612.1*) (Supplemental_Fig_S7A and Supplemental_Fig_S7B). Beyond testis, *AC1484477.2* nested gene high expression in two parts of the brain was associated with the expression of an increased number of isoforms of its host *GLANT9* (Figure 6C) whereas *AL117382.2* nested gene exclusive expression in the liver was associated with the expression of an isoform of *HNF4A* exclusively in this tissue (ENST00000372920.1, Supplemental_Fig_S8A). Given the correlation between co-expression and isoform regulation observed here and in previous studies (Cowley et al. 2012; Wood et al. 2008; Amante et al. 2020), we hypothesised that host genes would be more transcript diverse. Evaluation of isoform enrichment identified that in comparison to the transcriptome, host genes are isoform rich, in the GENCODE annotation (p<0.05, two-sample Kolmogorov-Smirnov test) (Figure 6B). We observed that host genes were enriched for longer genes (Supplemental_Fig_S1A), and because of the slight positive correlation between gene size and number of isoforms (Supplemental_Fig_S8B), we asked if the enrichment for higher number of isoforms was associated with host genes being longer. However, when we classified genes into size categories, we still observed that host genes were exhibiting a higher number of isoforms, suggesting that this was independent of their length (Supplemental_Fig_S8B).

## DISCUSSION

Global compilation of host/nested genes pairs in mammals illustrates their transcriptional profile during tissue-specification and interplay with RNA processing mechanisms and isoform regulation. Compared to the most exhaustive previous known list (Yu et al. 2005), we identified ∼50 times more pairs likely due to the continuous improvement of sequencing technologies allowing the annotation of more genes and transcripts. For example, between 2003 and 2013, the number of transcripts annotated in the RefSeq database increased by 23% (Pruitt et al. 2014). In addition, the comprehensive GENCODE annotation provides an exhaustive annotation for transcripts (Frankish et al. 2019, 2015). The resulting collection is the most comprehensive list of host/nested gene pairs in mouse and human which can be explored through the web application developed for this study (https://hngeneviewer.sites.er.kcl.ac.uk/hn_viewer/). Ultimately, the accuracy of the pair identification is dependent the genome annotation itself. The development of new technologies, such as long-read sequencing which will iteratively advance the detection of all possible genomic transcripts (Leung et al. 2021).

Contrary to the previous studies, a lower conservation of the host/nested genes was observed between mouse and human (between 21 and 34% of the pairs with known orthologues). This suggests that these events are mainly species-specific and the difference to previous studies can be explained by the updated annotation and inclusion of more diverse transcripts type (multiple types *versus* protein coding only). Furthermore, the conservation analysis is highly dependent on the annotation of the orthologue gene list, and it cannot be excluded that some conserved pairs were missed or excluded due to annotation defaults. A more detailed conservation study, including other species could improve the conservation level of host nested genes pairs. This would also be useful for investigating the origins of host/nested gene pairs and identify mechanisms involved such as *de novo* promoter generation inside a gene, transfer of a gene to inside another through genomic rearrangement or (retro)transposition, or extension of the host gene by acquisition of new exons which internalise an adjacent gene (Wright et al. 2021). These data indicate that most of the nested genes are fully contained within large introns of their hosts which suggests that there is a bias toward the acquisition of nested genes in these regions or a selection against events that interfere with the coding sequences of the host gene. This is also an indication that the impact of such a configuration would mainly be at the transcriptional level.

To test transcriptional impact of one gene on the other, we conducted a correlation analysis between host and nested gene expression across multiple tissues. Previous analysis suggested gene pairs were anti-correlated or that there was no relationship regarding expression (Assis et al. 2008; Yu et al. 2005). Our results agree with the latter with a lack of direct correlation, or anti-correlation between host and nested genes expression for most of the pairs. We identified that, only when co-expression is considered, the pairs show patterns related to tissue-specificity and not when partners, either host or nested, are examined on their own. This suggests that the organisation, pattern and potential function of the host and nested gene pairs expression could impact developmental processes and tissue specification. A high degree of co-expression of host and nested genes was observed in testis which correlates with reports of widespread transcription in this tissue (Melé et al. 2015). Because testis is a complex tissue, we leveraged single-cell RNA sequencing data where the different somatic and germ cell types are identifiable. In addition, we developed a novel analysis method to assess true co-expression in an individual cell-dependent manner. Thus, we investigated co-expression and mutual exclusive behaviour of host/nested genes in individual cells and showed that host and nested genes can be co-expressed in the same cells. We expect that the co-expression of pairs detected in single cells is an underestimate, given the limited depth of single cell sequencing and the poly(A) selection associated with most RNA-sequencing datasets. For most of the pairs the profile was complex and for some of them such as *IGSF11/IGSF11-AS1* changes in isoform diversity at the host is associated with expression of the nested gene only in testis. Previous work has demonstrated that the testis exhibits widespread gene activity alongside a higher transcriptome diversity compared to other tissues (Soumillon et al. 2013). We propose that one mechanism to provide this observed diversity, is through the widespread activity of nested genes that subsequently impacts isoform regulation of their corresponding host.

Even if more common in testis, we also observed a correlation between nested gene tissue-specific expression and host isoforms regulation in brain and liver. More globally, host genes were found to exhibit more transcript isoforms than other genes, suggesting an important role for the host/nested genes’ genomic organisation in the regulation of transcripts diversity. Previous works indicates that this can happened through the process of alternative polyadenylation (Cowley et al. 2012; Wood et al. 2008; Kaer et al. 2011), but the other mechanisms leading to transcript diversity, such as the use of alternative promoters or alternative splicing could be also regulated by the interplay between host and nested genes. The dissection of mechanisms involved would require the improvement of technical approaches, such as long read sequencing, capturing precisely the transcription start site, end site and splicing profile of individual RNA molecules at a sufficient depth.

## METHODS

### Identification of Host Nested Gene pairs in human (hg19) and mouse (mm10) genomes

To generate a list of all host/nested genes, comprehensive GENCODE annotations of the human (hg19, Release 36 (GRCh38)) and mouse (mm10, Release M25 (GRCm38.p6)) genomes were downloaded as a bed file from the UCSC table browser (https://genome.ucsc.edu/cgi-bin/hgTables). Transcripts denoted as, to be experimentally confirmed (TEC), immunoglobulin (IG) variable chain and T-cell receptor (TR) genes as well as complex loci (Protocadherin’s and UDP-glucuronosyltransferase) were removed from the list. The resulting list was used as a reference for transcripts for the hosts. To only select pairs where all the transcripts of the nested gene were included inside the host, the filtered list was also used to generate a ‘metagene’ list consisting of the most extreme coordinates of all possible transcripts of a single gene. Bedtools 2.29.2(Quinlan and Hall 2010) intersect function was used to overlap the metagenes list with reference list of transcripts for the hosts. The option –f 1.0 was used, meaning that the minimum overlap required was 100% of a metagene overlapping with another transcript. To keep only the overlap involving different genes, using an in-house R script genes which contained, the same name, the same size and where the start or end were the same were removed. When multiple transcripts from the same gene were involved in the same pair of genes, the longest transcript was kept. Annotation of opposite and same strand oriented pairs was determined by an in-house R script, checking if the strands of the pairs were: same (+:+ / -:-), or, opposite (+:- / -:+). A full list of host and nested genes is available in Supplemental_Table_S1 and Supplemental_Table_S2.

### Characterisation of the host/nested genes pairs

Information about transcripts biotypes, strand, number of isoforms and gene size was retrieved from GTF files downloaded from GENCODE website (https://www.gencodegenes.org/, Release M25 (GRCm38.p6) for mouse and Release 36 (GRCh37) for human). The functional enrichment analysis (Gene Ontology) was performed using g:Profiler (version e109_eg56_p17_1d3191d) with Bonferroni correction method applying significance threshold of 0.05 (Raudvere et al. 2019). The complete results are available in Supplemental_Table_S4. Data presented on host/nested gene pairs is viewable via https://hngeneviewer.sites.er.kcl.ac.uk/hn_viewer/ which was developed using RShiny.

### Annotation of the nested gene location

ChIPseeker (version 1.24.0)(Wang et al. 2022) was used to annotate the location of the nested genes inside their host. A custom GTF file containing only the information about the host transcripts involved in host/nested genes pairs was used as a reference and a bed file containing the start and end coordinates of the nested genes was used as a query. The genomicAnnotationPriority was define as “Exon”,“Intron”,“Promoter”,“5UTR”,“3UTR”,“Downstream” and “Intergenic”. The pairs with successful nested gene annotation inside their host were retained and further rounds of annotation used when necessary (after removal of the already annotated pairs) to annotate the missing pairs (for example when a gene is nested inside two different genes). Finally, for the last pairs with missing annotation, manual annotation was performed. The table containing the annotation and exon/intron numbers was retrieved and cross referenced with the GENCODE GTF file for intron/exon characteristics.

### Conservation analysis

The list of all the human-mouse orthologs was downloaded using the Ensembl BioMart software suite (Cunningham et al. 2022) with the following filter “Homolog filters” option “Orthologous Mouse Genes” from the human gene list. The table was next downloaded with the following attributes: Gene stable ID, Transcript stable ID, Mouse gene stable ID, Mouse gene name, Mouse protein or transcript stable ID. The orthologs information was next overlapped with the lists of host/nested genes using an in-house R script to identify the 349 combinations of orthologous pairs (Supplemental_Table_S3).

### ENCODE RNA-Sequencing analysis

Paired end RNA sequencing data from multiple tissues was used in both human and mouse (for accession numbers, see Sup Tables 4,5). Reads were aligned using kallisto (Bray et al. 2016) and a read count table per sample was generated using tximport (Soneson et al. 2016); reads were then normalised using DESeq2 (Love et al. 2014). Pairs of host/nested genes were processed with a pair_id which was used to sort normalised read matrix data based on the corresponding host and nested gene ENST_ID using a custom R script. Correlations of host/nested gene expression across all samples was performed using the Spearman’s rank correlation coefficient. Highly correlated (coefficient >0.7) host/nested gene pairs were hierarchically clustered using the complete-linkage clustering method using the pheatmap package. Tissue-specificity of host, nested and all genes were evaluated using standard deviation of expression across all tissues, and, calculation of Tau index (Yanai et al. 2005) using the package tspex (Camargo et al. 2020) on the normalised count matrix.

### scRNA-sequencing analysis

A table with Integrated, normalized counts per cell from scRNA-seq dataset during spermatogenesis was retrieved from Gene Expression Omnibus (GEO) database (accession number GSE153947) (Di Persio et al. 2021). The dataset was filtered to only include host and nested genes which were highly correlated and expressed in testis from the ENCODE-RNA sequencing analysis. The expression of a gene was reduced to a binary signal whereby if expression reached above the threshold (a quarter of the maximum expression), a value of 1 was given, and a value of 0 if below this threshold. Host/nested genes pairs were co-expressed in a single cell if the sum of this value was 2. If the sum equal to 1, then single cells were only expressing either the host or nested gene. If the sum equaled 0, the single cells were not expressing either gene. A schematic illustrating this method is available in Supplemental_Fig_3B. For further analysis, pairs where over 1% of cells were expressing both transcripts were considered. The proportion of cells co-expressing each host/nested gene pairs was clustering via k-means clustering using the factoextra package (Version 1.0.7.999).

## Supporting information

All Supplementary Figures with Legends

## ACKNOWLEDGEMENTS

We thank Hannah Mischo for advice and useful discussions. We thank Lukasz Zalewski and Stuart Morrison for their assistance with the R Shiny app. We thank Hallgerdur Kolbeinsdottir for critical reading of the manuscript. The Genotype-Tissue Expression (GTEx) Project was supported by the Common Fund of the Office of the Director of the National Institutes of Health, and by NCI, NHGRI, NHLBI, NIDA, NIMH, and NINDS. The data used for the analyses described in this manuscript were obtained from: the GTEx Portal on 13/02/24.

## AUTHOR CONTRIBUTIONS

BM and JC concieved the idea, performed the analysis, and wrote the manuscript. JC developed the shiny app with support and guidance from RTM. RJO provided supervision, discussed the work with BM and JC and critically reviewed and contributed to the final manuscript.

## REFERENCES

Amante SM, Montibus B, Cowley M, Barkas N, Setiadi J, Saadeh H, Giemza J, Contreras-Castillo S, Fleischanderl K, Schulz R, et al. 2020. Transcription of intragenic CpG islands influences spatiotemporal host gene pre-mRNA processing. Nucleic Acids Res 48: 8349–8359.

Assis R, Kondrashov AS, Koonin EV, Kondrashov FA. 2008. Nested genes and increasing organizational complexity of metazoan genomes. Trends Genet 24: 475–478.

Billingsley DJ, Bonass WA, Crampton N, Kirkham J, Thomson NH. 2012. Single-molecule studies of DNA transcription using atomic force microscopy. Phys Biol 9: 021001.

Bray NL, Pimentel H, Melsted P, Pachter L. 2016. Near-optimal probabilistic RNA-seq quantification. Nat Biotechnol 34: 525–527.

Camargo AP, Vasconcelos AA, Fiamenghi MB, Pereira GAG, Carazzolle MF. 2020. tspex: a tissue-specificity calculator for gene expression data. https://www.researchsquare.com (Accessed May 12, 2022).

Chen B, Zhu G, Yan A, He J, Liu Y, Li L, Yang X, Dong C, Kee K. 2021. IGSF11 is required for pericentric heterochromatin dissociation during meiotic diplotene. PLOS Genet 17: e1009778.

Chen C-H, Pan C-Y, Lin W. 2019. Overlapping protein-coding genes in human genome and their coincidental expression in tissues. Sci Rep 9: 13377.

Cowley M, Wood AJ, Böhm S, Schulz R, Oakey RJ. 2012. Epigenetic control of alternative mRNA processing at the imprinted Herc3/Nap1l5 locus. Nucleic Acids Res 40: 8917– 8926.

Cunningham F, Allen JE, Allen J, Alvarez-Jarreta J, Amode MR, Armean IM, Austine-Orimoloye O, Azov AG, Barnes I, Bennett R, et al. 2022. Ensembl 2022. Nucleic Acids Res 50: D988–D995.

Davis CA, Hitz BC, Sloan CA, Chan ET, Davidson JM, Gabdank I, Hilton JA, Jain K, Baymuradov UK, Narayanan AK, et al. 2018. The Encyclopedia of DNA elements (ENCODE): data portal update. Nucleic Acids Res 46: D794–D801.

Di Persio S, Tekath T, Siebert-Kuss LM, Cremers J-F, Wistuba J, Li X, Meyer zu Hörste G, Drexler HCA, Wyrwoll MJ, Tüttelmann F, et al. 2021. Single-cell RNA-seq unravels alterations of the human spermatogonial stem cell compartment in patients with impaired spermatogenesis. Cell Rep Med 2: 100395.

Dowd C. 2020. A New ECDF Two-Sample Test Statistic. http://arxiv.org/abs/2007.01360 (Accessed October 7, 2022).

Feiss M, Fisher RA, Crayton MA, Egner C. 1977. Packaging of the bacteriophage λ chromosome: Effect of chromosome length. Virology 77: 281–293.

Frankish A, Diekhans M, Ferreira A-M, Johnson R, Jungreis I, Loveland J, Mudge JM, Sisu C, Wright J, Armstrong J, et al. 2019. GENCODE reference annotation for the human and mouse genomes. Nucleic Acids Res 47: D766–D773.

Frankish A, Uszczynska B, Ritchie GR, Gonzalez JM, Pervouchine D, Petryszak R, Mudge JM, Fonseca N, Brazma A, Guigo R, et al. 2015. Comparison of GENCODE and RefSeq gene annotation and the impact of reference geneset on variant effect prediction. BMC Genomics 16: S2.

Henikoff S, Keene MA, Fechtel K, Fristrom JW. 1986. Gene within a gene: nested Drosophila genes encode unrelated proteins on opposite DNA strands. Cell 44: 33–42.

Jan SZ, Vormer TL, Jongejan A, Röling MD, Silber SJ, de Rooij DG, Hamer G, Repping S, van Pelt AMM. 2017. Unraveling transcriptome dynamics in human spermatogenesis. Dev Camb Engl 144: 3659–3673.

Jia Z, Wu Q. 2020. Clustered Protocadherins Emerge as Novel Susceptibility Loci for Mental Disorders. Front Neurosci 14. https://www.frontiersin.org/articles/10.3389/fnins.2020.587819/full (Accessed February 19, 2021).

Jung D, Giallourakis C, Mostoslavsky R, Alt FW. 2006. Mechanism and control of V(D)J recombination at the immunoglobulin heavy chain locus. Annu Rev Immunol 24: 541– 570.

Kaer K, Branovets J, Hallikma A, Nigumann P, Speek M. 2011. Intronic L1 Retrotransposons and Nested Genes Cause Transcriptional Interference by Inducing Intron Retention, Exonization and Cryptic Polyadenylation. PLOS ONE 6: e26099.

Latos PA, Pauler FM, Koerner MV, Şenergin HB, Hudson QJ, Stocsits RR, Allhoff W, Stricker SH, Klement RM, Warczok KE, et al. 2012. Airn Transcriptional Overlap, But Not Its lncRNA Products, Induces Imprinted Igf2r Silencing. Science 338: 1469–1472.

Leung SK, Jeffries AR, Castanho I, Jordan BT, Moore K, Davies JP, Dempster EL, Bray NJ, O’Neill P, Tseng E, et al. 2021. Full-length transcript sequencing of human and mouse cerebral cortex identifies widespread isoform diversity and alternative splicing. Cell Rep 37. https://www.cell.com/cell-reports/abstract/S2211-1247(21)01504-7 (Accessed November 17, 2021).

Li WV, Li Y. 2021. scLink: Inferring Sparse Gene Co-expression Networks from Single-cell Expression Data. Genomics Proteomics Bioinformatics 19: 475–492.

Licatalosi DD, Darnell RB. 2010. RNA processing and its regulation: global insights into biological networks. Nat Rev Genet 11: 75–87.

Lonsdale J, Thomas J, Salvatore M, Phillips R, Lo E, Shad S, Hasz R, Walters G, Garcia F, Young N, et al. 2013. The Genotype-Tissue Expression (GTEx) project. Nat Genet 45: 580–585.

Love MI, Huber W, Anders S. 2014. Moderated estimation of fold change and dispersion for RNA-seq data with DESeq2. Genome Biol 15: 550.

Melé M, Ferreira PG, Reverter F, DeLuca DS, Monlong J, Sammeth M, Young TR, Goldmann JM, Pervouchine DD, Sullivan TJ, et al. 2015. The human transcriptome across tissues and individuals. Science 348: 660–665.

Neri F, Rapelli S, Krepelova A, Incarnato D, Parlato C, Basile G, Maldotti M, Anselmi F, Oliviero S. 2017. Intragenic DNA methylation prevents spurious transcription initiation. Nature 543: 72–77.

Pollen AA, Nowakowski TJ, Shuga J, Wang X, Leyrat AA, Lui JH, Li N, Szpankowski L, Fowler B, Chen P, et al. 2014. Low-coverage single-cell mRNA sequencing reveals cellular heterogeneity and activated signaling pathways in developing cerebral cortex. Nat Biotechnol 32: 1053–1058.

Prescott EM, Proudfoot NJ. 2002. Transcriptional collision between convergent genes in budding yeast. Proc Natl Acad Sci 99: 8796–8801.

Pruitt KD, Brown GR, Hiatt SM, Thibaud-Nissen F, Astashyn A, Ermolaeva O, Farrell CM, Hart J, Landrum MJ, McGarvey KM, et al. 2014. RefSeq: an update on mammalian reference sequences. Nucleic Acids Res 42: D756–D763.

Quinlan AR, Hall IM. 2010. BEDTools: A flexible suite of utilities for comparing genomic features. Bioinformatics 26: 841–842.

Raudvere U, Kolberg L, Kuzmin I, Arak T, Adler P, Peterson H, Vilo J. 2019. g:Profiler: a web server for functional enrichment analysis and conversions of gene lists (2019 update). Nucleic Acids Res 47: W191–W198.

Sanchez-Taltavull D, Perkins TJ, Dommann N, Melin N, Keogh A, Candinas D, Stroka D, Beldi G. 2020.Bayesian correlation is a robust gene similarity measure for single-cell RNA-seq data. NAR Genomics Bioinforma 2: lqaa002.

Simon-Loriere E, Holmes EC, Pagán I. 2013. The Effect of Gene Overlapping on the Rate of RNA Virus Evolution. Mol Biol Evol 30: 1916–1928.

Singh I, Lee S-H, Sperling AS, Samur MK, Tai Y-T, Fulciniti M, Munshi NC, Mayr C, Leslie CS. 2018. Widespread intronic polyadenylation diversifies immune cell transcriptomes. Nat Commun 9: 1716.

Soneson C, Love MI, Robinson MD. 2016. Differential analyses for RNA-seq: transcript-level estimates improve gene-level inferences. https://f1000research.com/articles/4-1521 (Accessed October 5, 2022).

Soumillon M, Necsulea A, Weier M, Brawand D, Zhang X, Gu H, Barthès P, Kokkinaki M, Nef S, Gnirke A, et al. 2013. Cellular Source and Mechanisms of High Transcriptome Complexity in the Mammalian Testis. Cell Rep 3: 2179–2190.

Spencer CA, Gietz RD, Hodgetts RB. 1986. Overlapping transcription units in the dopa decarboxylase region of Drosophila. Nature 322: 279–281.

Strassburg CP, Kalthoff S, Ehmer U. 2008. Variability and Function of Family 1 Uridine-5′-Diphosphate Glucuronosyltransferases (UGT1A). Crit Rev Clin Lab Sci 45: 485–530.

Veeramachaneni V, Makalowski W, Galdzicki M, Sood R, Makalowska I. 2004. Mammalian Overlapping Genes: The Comparative Perspective. Genome Res 14: 280–286.

Wang Q, Li M, Wu T, Zhan L, Li L, Chen M, Xie W, Xie Z, Hu E, Xu S, et al. 2022.Exploring Epigenomic Datasets by ChIPseeker. Curr Protoc 2: e585.

Weisbeek PJ, Borrias WE, Langeveld SA, Baas PD, Arkel GAV. 1977. Bacteriophage phiX174: gene A overlaps gene B. Proc Natl Acad Sci 74: 2504–2508.

Williams T, Fried M. 1986. A mouse locus at which transcription from both DNA strands produces mRNAs complementary at their 3′ ends. Nature 322: 275–279.

Wood AJ, Schulz R, Woodfine K, Koltowska K, Beechey CV, Peters J, Bourc’his D, Oakey RJ. 2008. Regulation of alternative polyadenylation by genomic imprinting. Genes Dev 22: 1141–1146.

Wright BW, Molloy MP, Jaschke PR. 2021. Overlapping genes in natural and engineered genomes. Nat Rev Genet 1–15.

Wu Z, Yang H, Colosi P. 2010. Effect of Genome Size on AAV Vector Packaging. Mol Ther 18: 80–86.

Yanai I, Benjamin H, Shmoish M, Chalifa-Caspi V, Shklar M, Ophir R, Bar-Even A, Horn-Saban S, Safran M, Domany E, et al. 2005. Genome-wide midrange transcription profiles reveal expression level relationships in human tissue specification. Bioinformatics 21: 650–659.

Yu P, Ma D, Xu M. 2005. Nested genes in the human genome. Genomics 86: 414–422.

