## Supplementary material for "Global identification of mammalian host and nested gene pairs reveal tissue-specific transcriptional interplay": All Supplementary Figures with Legends

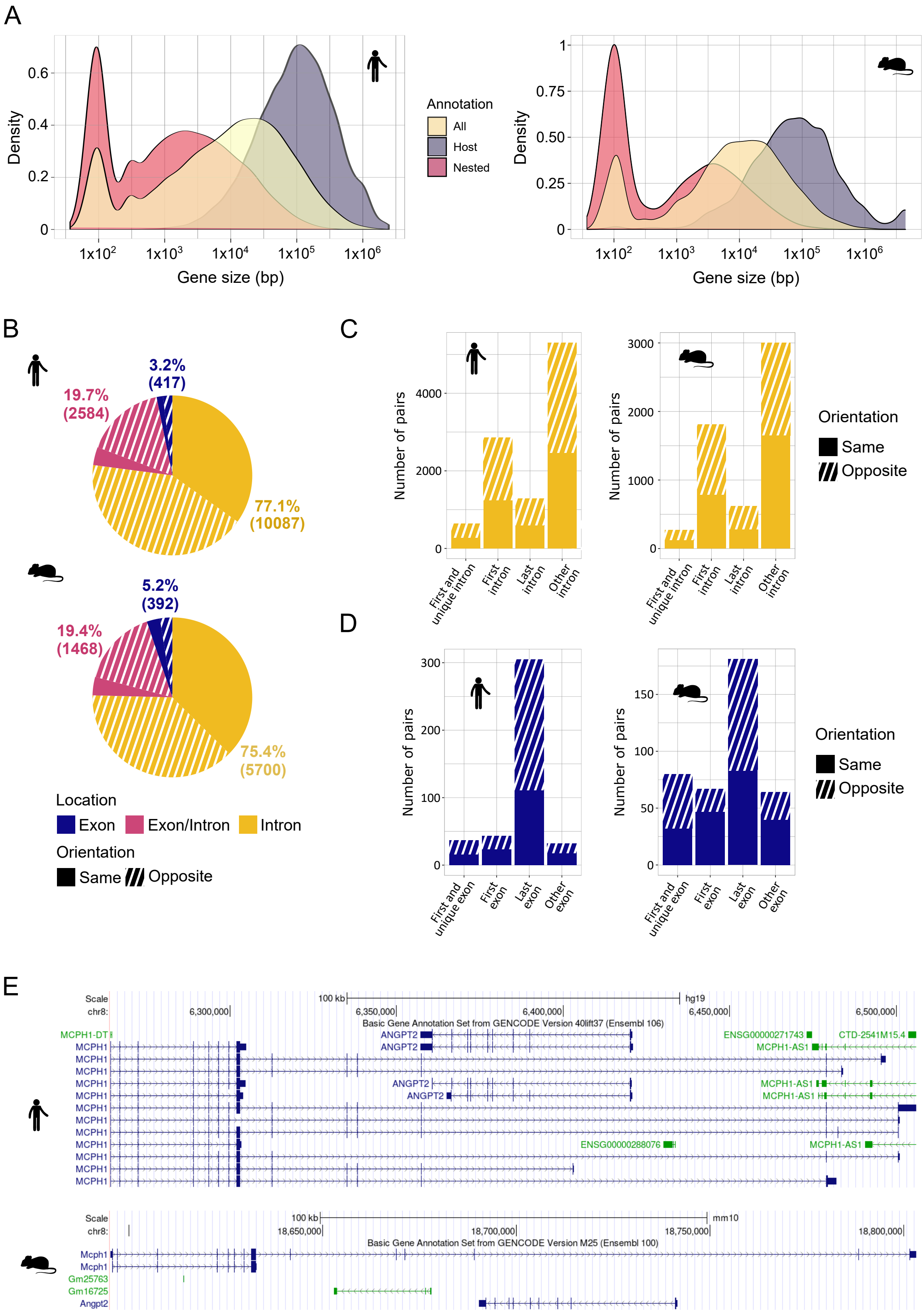
A
Density
0.6
0.4
0.2
0
1x10^2
1x10^3
1x10^4
1x10^5
1x10^6
Gene size (bp)
Annotation
All
Host
Nested
1
0.75
0.50
0.25
B
19.7%
(2584)
3.2%
(417)
77.1%
(10087)
19.4%
(1468)
5.2%
(392)
75.4%
(5700)
Location
Exon
Exon/Intron
Intron
Orientation
Same
Opposite
C
Number of pairs
4000
2000
3000
1000
First and unique intron
First intron
Last intron
Other intron
D
300
200
100
150
50
First and unique exon
First exon
Last exon
Other exon
E
Scale
chr8:
100 kb
hg19
6,300,000
6,350,000
6,400,000
6,450,000
6,500,000
Basic Gene Annotation Set from GENCODE Version 40lift37 (Ensembl 106)
MCPH1-DT
MCPH1
ANGPT2
ENSG00000271743
CTD-2541M15.4
MCPH1-AS1
ENSG00000288076
mm10
18,650,000
18,700,000
18,750,000
18,800,000
Basic Gene Annotation Set from GENCODE Version M25 (Ensembl 100)
Mcph1
Gm25763
Gm16725
Angpt2

**Supplemental_Fig_S1. Host/nested genes pairs in mouse and human shows similar characteristics.** (A) Distribution of the genes' size for host genes, nested genes, and all genes. A base-10 log scale was used for the gene size. (B) Distribution of the locations of nested genes within their host genes according to their orientation to each other. (C) Detailed location of the nested genes when fully contained inside an intron. (D) Detailed location of the nested genes when fully contained inside an exon. (E) Screenshots of the UCSC genome browser for an example of a host/nested genes pair conserved between mouse and human.

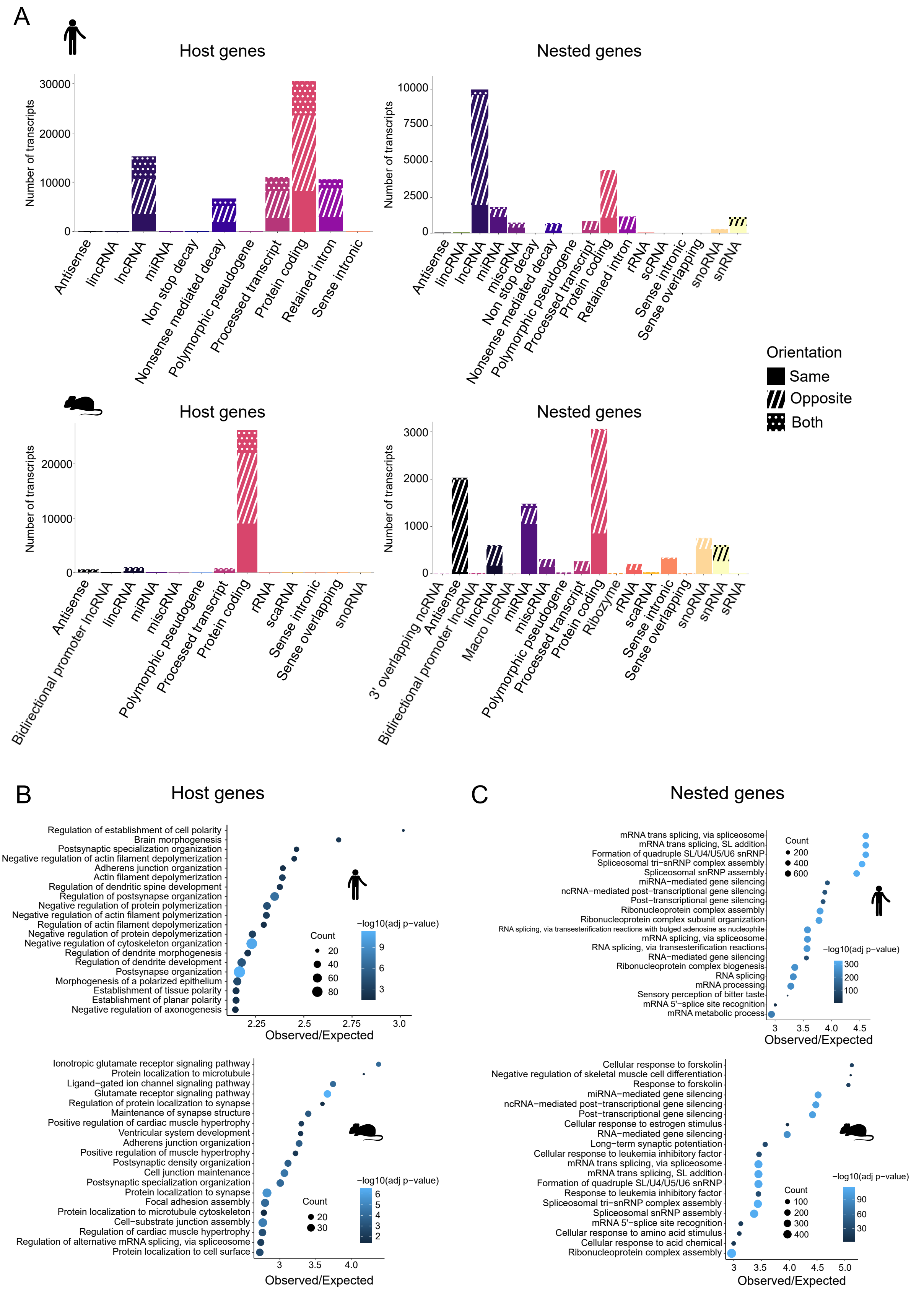


**Supplemental_Fig_S2. Host/nested genes pair's biotypes, functional enrichment analysis and expression dispersion.** (A) Number of host and nested genes associated with each biotype according to their orientation in pairs. Both refers to genes involved in multiple pairs with different relative orientation. (B) Gene sets enrichment analysis for host gene lists. Only the top 20 (according to the observed/expected ratio) significant biological processes are represented. (C) Gene sets enrichment analysis for nested gene lists. Only the top 20 (according to the observed/expected ratio) significant biological processes are represented.

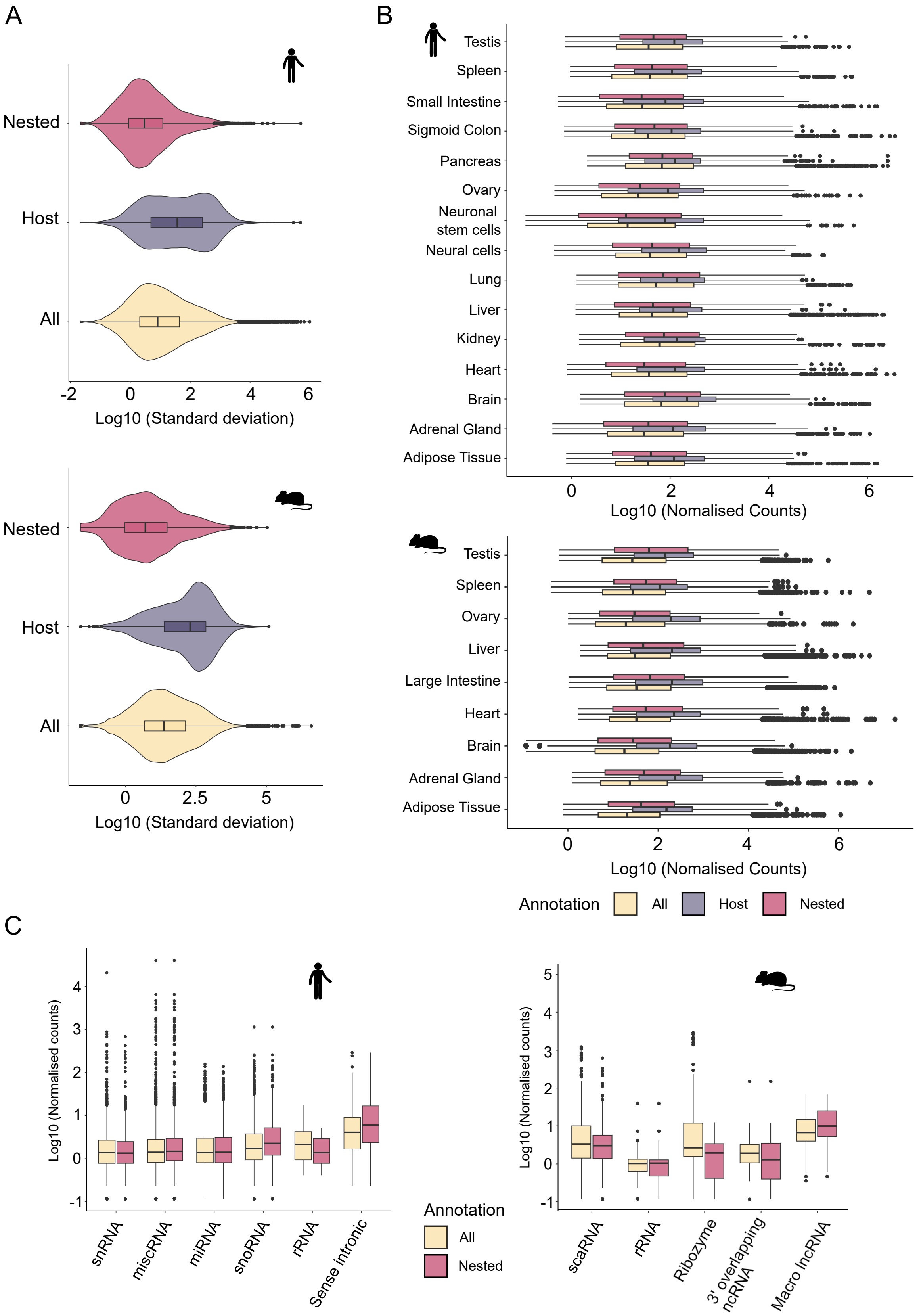


**Supplemental_Fig_S3. Distribution of the expression of host/nested genes in ENCODE RNA-sequencing datasets.** (A) Distribution of standard deviation between the normalised counts across tissues (B) Distribution of the expression of all, host and nested genes for protein coding genes across tissues, in human and mouse transcriptomes (C) Distribution of the expression of all and nested genes for biotypes unique to nested genes, in human and mouse transcriptomes

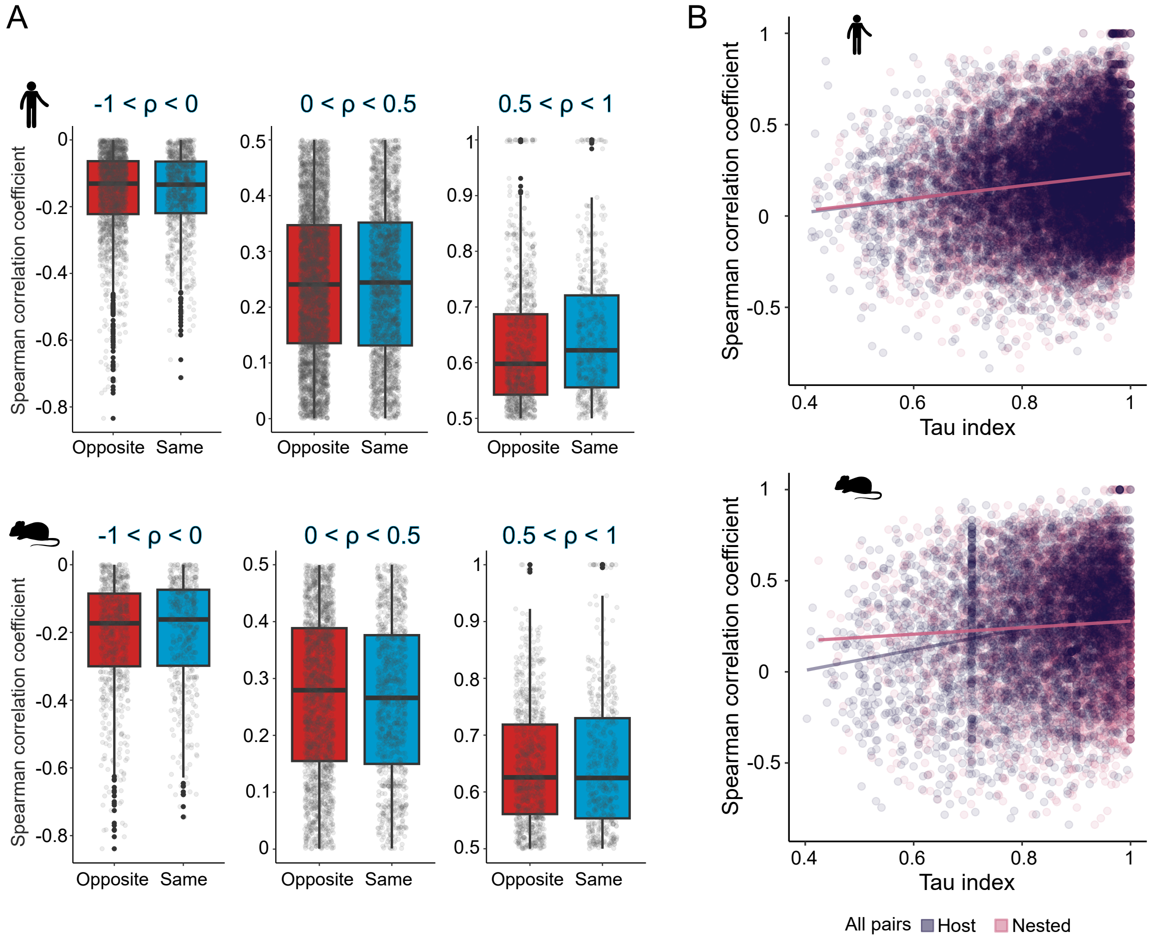


**Supplemental_Fig_S4. Expression correlation between host and nested genes and its association with tissue specificity.** (A) Distribution of Spearman's rank correlation coefficient between pairs according to their orientation and depending on the range of ρ. (B) Correlation between the τ index and Spearman's rank correlation coefficient across all host nested gene pairs.

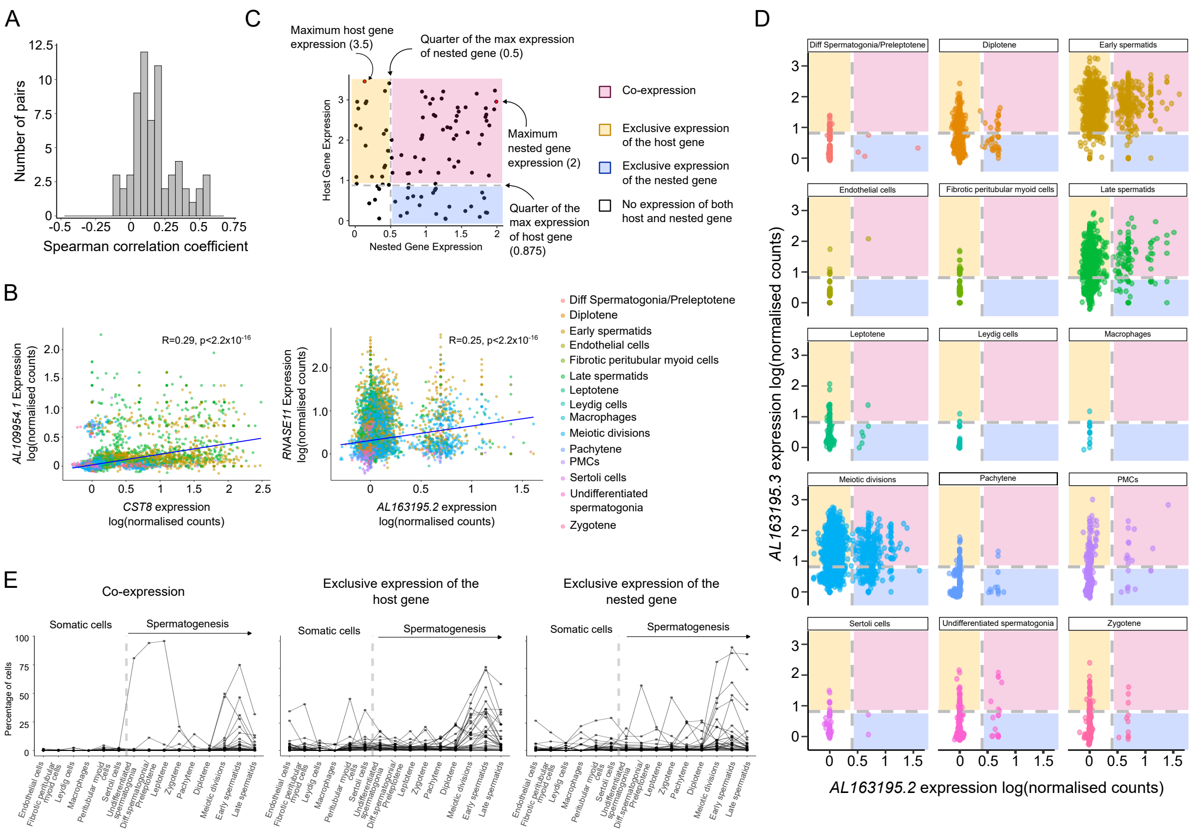


**Supplemental_Fig_S5. Host and nested genes with high co-expression in testis can exhibit dynamic co-expression patterns during spermatogenesis.** (A) Distribution of the Spearman's rank correlation coefficient values between expression of the host and nested genes across testis single cells (B) Scatter plots showing examples of host and nested gene expression and Spearman's rank correlation across single cells in testis for *AL109954.1/CST8*, R=0.29 and *RNASE11/AL163195.2* R= 0.25 (C) Schematic representation of the filtering methodology applied to single cells to determine proportion of cells which co-express a host and nested gene pair, or only express one of the partner. In this randomised mock example, host (0.875) and nested (0.5) gene expression thresholds were used to calculate that 46% of single cells co-express the gene pair. (D) Scatter plots showing examples of host and nested genes co-expression in the different testis cell types for the pair *AL163195.3/AL163195.2* (E) Percentage of cells co-expressing host and nested gene pairs or expressing only one of the partners across spermatogenesis for the 34 pairs with tissue specific expression in testis and robust expression data in the scRNA-seq.

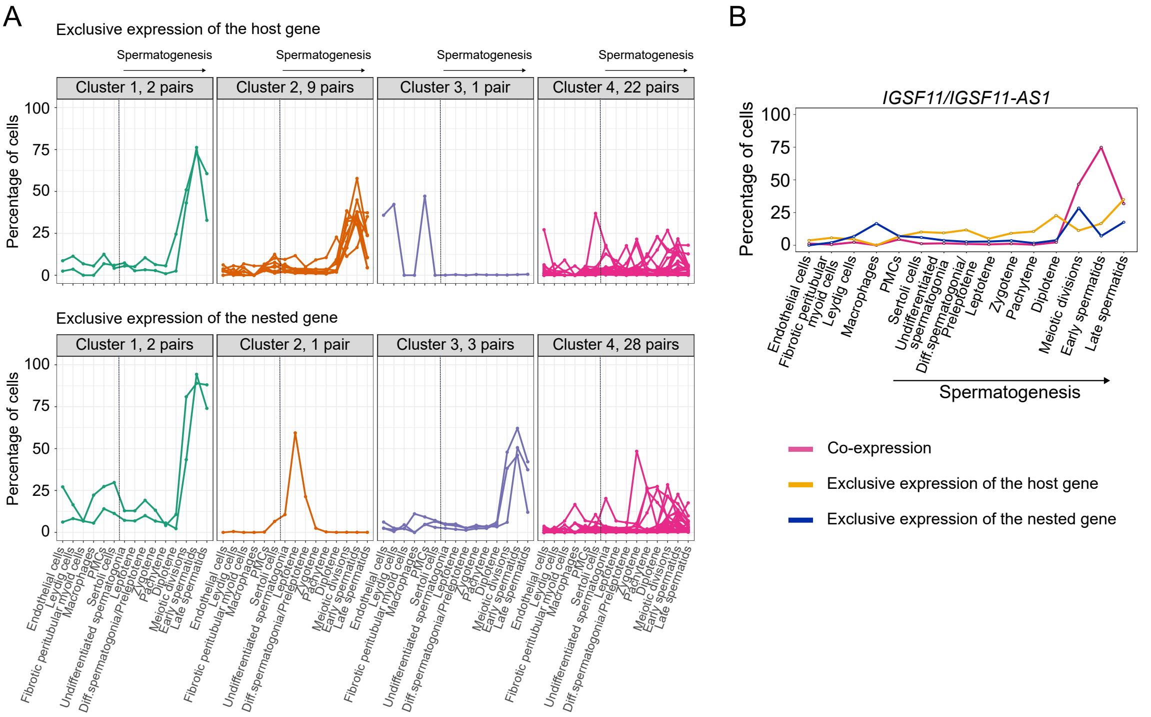


**Supplemental_Fig_S6. Profiles of host and nested genes expression during spermatogenesis.** (A) Proportion of cells which only express host or nested gene across spermatogenesis. Four distinct co-expression profiles were determined by K-means clustering. (B) Scatter plot showing the profiles of expression determined as in Supplementary Fig5C for *IGSF11/IGSF11-AS1*.

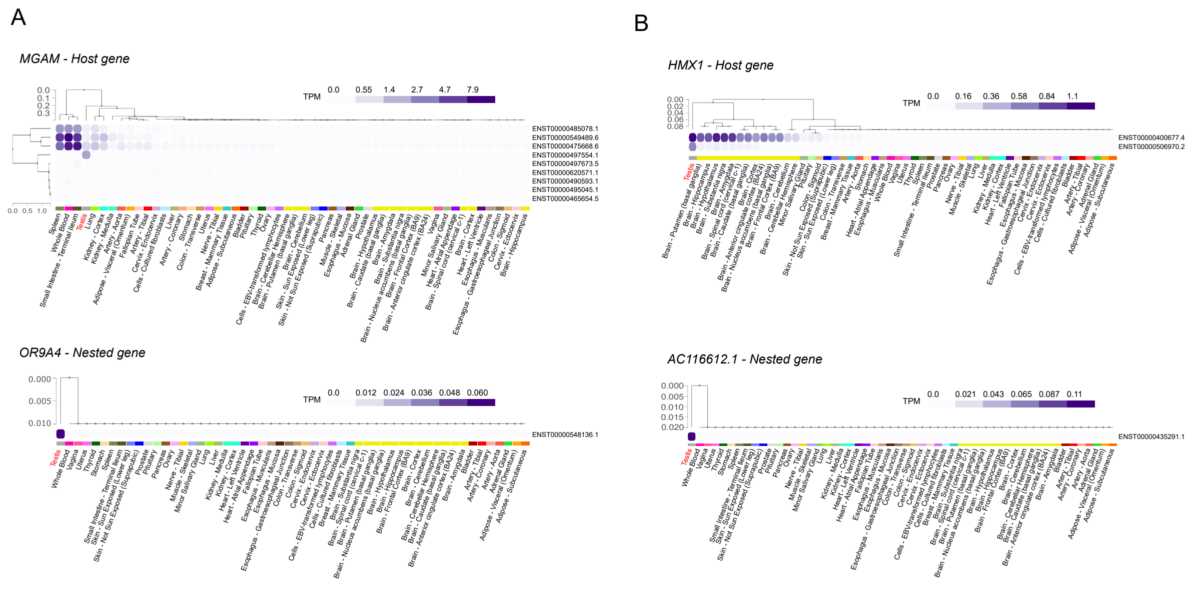


**Supplemental_Fig_S7. Host and nested genes co-expression is correlated with regulation of isoform diversity.** (A) Profile of expression of the different isoforms (rows) of the *MGAM/OR9A4* pair in different tissues (columns) from the GTEx portal data on isoforms expression (https://gtexportal.org/home/). Isoform and tissue were ordered by hierarchal clustering using the Euclidean distance and average linkages. Tissues where co-expression is happening are in red. (B) Profile of expression of the different isoforms (rows) of the *HMX1/AC116612.1* pair in different tissues (columns) from the GTEx portal data on isoforms expression (https://gtexportal.org/home/). Isoform and tissue were ordered by hierarchal clustering using the Euclidean distance and average linkages. Tissues where co-expression is happening are in red.

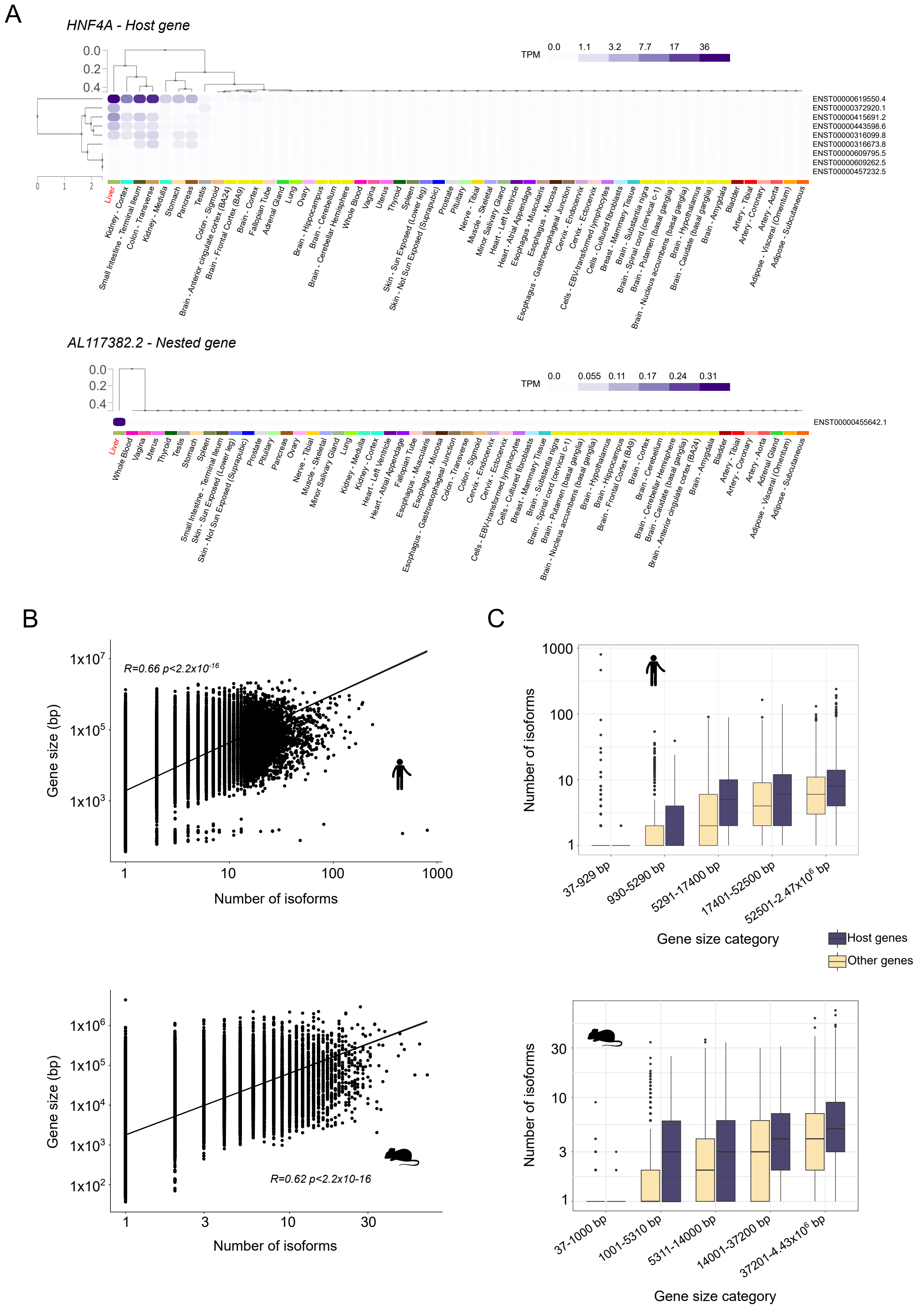


**Supplemental_Fig_S8**. **Host genes display a higher number of isoforms.** (A) Profile of expression of the different isoforms (rows) of the *HNF4A/AL117382.2* pair in different tissues (columns) from the GTEx portal data on isoforms expression (https://gtexportal.org/home/). Isoform and tissue were ordered by hierarchal clustering using the Euclidean distance and average linkages. Tissues where co-expression is happening are in red. (B) Scatter plot showing the correspondence between the gene size and number of isoforms. A base-10 log scale was used for both axes. The correlation was determined by Spearman's rank correlation. (C) Distribution of the number of isoforms per size categories. Genes were classified according to the size of the longest isoform.  Each bin contains an equal total number of genes.

**Supplemental_Table_S1. List and characteristics of host and nested genes pairs in human.**

**Supplemental_Table_S2. List and characteristics of host and nested genes pairs in mouse.**

**Supplemental_Table_S3. Conserved host/nested gene pairs between human and mouse genomes.**

**Supplemental_Table_S4. GO analysis on host and nested genes lists.**

**Supplemental_Table_S5. ENCODE metadata human**

**Supplemental_Table_S6. ENCODE metadata mouse**
